# Specific deletion of interleukin-1 beta in microglia improves acute outcome and modulates neurogenesis after ischemic stroke

**DOI:** 10.64898/2025.12.19.695391

**Authors:** Alba Grayston, Margarida Baptista, Kelly Wemyss, Ruby Taylor, Grace Cullen, Syeda S Jafree, Nadim Luka, Joshua R Cox, Joanne E Konkel, David Brough, Stuart M Allan, Emmanuel Pinteaux

**Affiliations:** Division of Neuroscience, School of Biological Sciences, Faculty of Biology, Medicine and Health (FBMH), The University of Manchester, Manchester, UK; Geoffrey Jefferson Brain Research Centre, University of Manchester, Northern Care Alliance NHS Foundation Trust, The Manchester Academic Health Science Centre, Manchester, UK; Lydia Becker Institute of Immunology and Inflammation, School of Biological Sciences, Faculty of Biology, Medicine and Health, Manchester Academic Health Science Centre, University of Manchester, Manchester, UK

**Keywords:** ischemic stroke, microglia, interleukin-1 beta

## Abstract

Interleukin-1 (IL-1) signaling is a major driver of post-ischemic neuroinflammation, yet the cell- and isoform-specific roles of the two major IL-1 receptor type 1 agonists, IL-1α and IL-1β, remain incompletely defined in the context of stroke. Microglia rapidly express IL-1α after cerebral ischemia, whereas IL-1β expression is delayed and restricted to a small subset of microglia and infiltrating immune cells. Here, we investigated for the first time the specific contribution of microglial-derived IL-1β to acute injury and post-stroke neurorepair after transient middle cerebral artery occlusion in male and female mice, through microglial-specific tamoxifen-inducible Cre-loxP-mediated recombination. Deletion of microglial IL-1β improved acute neurological outcome, reduced neutrophil accumulation in the ischemic brain and dampened systemic inflammatory cytokines. These effects were most evident during the acute phase and in female in mice. In contrast, long-term functional recovery was largely unaffected. However, microglial IL-1β deletion differentially regulated post-stroke neurogenesis, enhancing subventricular zone neurogenic responses and ectopic neuroblast migration while limiting hippocampal neurogenesis. Together, these findings identify microglial IL-1β as a key amplifier of early inflammatory injury after stroke, exerting region-specific effects on neurogenic niches, and highlight distinct, non-redundant roles for microglial IL-1 isoforms in ischemic brain injury and repair.

## 1. Introduction

Interleukin-1 (IL-1) is a central driver of the inflammatory response after ischemic stroke. Extensive pre-clinical and clinical evidence demonstrates that IL-1 exacerbates ischemic injury, while blocking IL-1 actions using the IL-1 receptor type 1 (IL-1R1) antagonist (IL-1Ra) improves outcomes in experimental stroke^1^. However, the cellular sources and isoform-specific actions of IL-1 in the post-stroke brain remain incompletely understood. IL-1 is highly pleiotropic, and the two main IL-1R1 agonists, IL-1α and IL-1β, though they bind the same receptor and elicit the same cellular downstream pathways, exert distinct non-redundant functions during sterile and infectious inflammation and show unique regulation at the organ/tissue level, release mechanisms, and cellular targets^2^. Furthermore, it has been shown that the consequences of IL-1 signaling on immunological, neural, and physiological activities are controlled by discrete cell-type-specific IL-1R1 systems^3^. Understanding the cellular source and targets, spatiotemporal expression and functional consequences of the IL-1 family are essential to guide the development of selective IL-1 targeting therapies for brain injury. In the context of ischemic stroke, microglia are key early drivers of post-stroke neuroinflammation^4^. Previous work from our group has shown that IL-1α is rapidly and robustly induced in microglia within hours of cerebral ischemia, whereas IL-1β follows a delayed expression in a small subset of microglia and in infiltrating peripheral immune cells^5, 6^. Our recent study suggested that microglial-derived IL-1α is dispensable for acute stroke injury but is required for neurorepair processes and long-term functional recovery, highlighting isoform- and cell-specific functions of IL-1 signaling after stroke^6^. Although IL-1β is widely recognized as a key inflammatory cytokine released during sterile brain injury and has been implicated in the regulation of neurogenesis^7, 8^, the specific contribution of microglial-derived IL-1β to stroke pathogenesis and recovery is largely unknown. Importantly, few studies have interrogated IL-1 biology in both sexes, despite evidence that inflammatory networks and stroke outcomes differ between males and females. Here, building on our previous work dissecting the role of microglial IL-1α, we investigated the specific contribution of microglial-derived IL-1β to acute and long-term stroke outcome. Using a genetic model enabling selective deletion of IL-1β in microglia as the sole resident brain cell population that express IL-1β following cerebral ischemia, we assessed the impact of microglial IL-1β deletion on immune responses, neuroinflammation, neurogenesis, angiogenesis and functional recovery after stroke, in both male and female mice. This approach allowed us to determine whether microglial IL-1β plays a role distinct from, or complementary to, microglial IL-1α in shaping injury and repair after ischemic stroke and if there is sex dependency to any effects observed.

## 2. Methods

### 2.1. Animals

All animal procedures were carried out in accordance with the Animals (Scientific Procedures) Act (1986), under a Home Office UK project license, approved by the local Animal Welfare Ethical Review Board, and experiments were performed in accordance with ARRIVE (Animal Research: Reporting of In Vivo Experiments) guidelines^9^, with researchers blinded to genotype.

Animals were housed at 21±1°C, 55±10% humidity on a 12h light-dark cycle in Sealsafe Plus Mouse individually ventilated cages (Techniplast, Italy). Mice were supplied with Sizzle Nest nesting material (Datesand Ltd., UK) and cardboard enrichment tubes (Datesand Ltd., UK), had *ad libitum* access to standard rodent diet (SDS, UK) and water.

Experiments were performed on a total of 114 C57BL/6J mice (46 female and 68 male) from an in-house colony (IL-1β^fl/fl^:Cx3cr1-Cre^ERT2^ and IL-1β^fl/fl^ as littermate controls) at the University of Manchester. Complete schemes of experimental designs are presented in Figures 1-5. A total of 15 mice were excluded: 1 due to experimental failure/complications, 1 due to mortality post-intervention and 13 culled due to declining post-stroke health issues and in line with regulatory requirements. Microglial specific IL-1β knockout (KO) mice were generated by crossing IL-1β^fl/fl^ mice in which exons 4 and 5 of the *IL1B* gene is flanked with loxP sites ^10^ with Cx3Cr1-Cre^ERT2^ mice expressing CX3CR1 promoter-driven Cre recombinase (JAX stock #020940) ^11^ that is expressed in the mononuclear phagocyte lineage. To induce Cre^ERT2^ activity and subsequent Cre-loxP-mediated deletion of IL-1β in CX3CR1-expressing cells, 3-6 months old mice were given tamoxifen by intraperitoneal injection for 5 consecutive days (2mg/100µL in corn oil, 75 mg/kg, Sigma-Aldrich). To exclude potential effects from a partial KO of the fractalkine receptor, CX3CR1, IL-1β^fl/fl^:Cx3cr1-Cre^ERT2^ mice without tamoxifen treatment (wild type, WT Cre+) were included alongside Cre^-/-^ IL-1β^fl/fl^ treated with tamoxifen (WT Cre-). To reduce animal numbers, only WT Cre-mice were used as a control group in the absence of a partial CX3CR1 KO effect. Mice were recovered for 4 weeks after tamoxifen treatment to allow the peripheral myeloid WT population to replenish, with sustained IL-1β deletion in microglia^10^.

**Figure 1.**
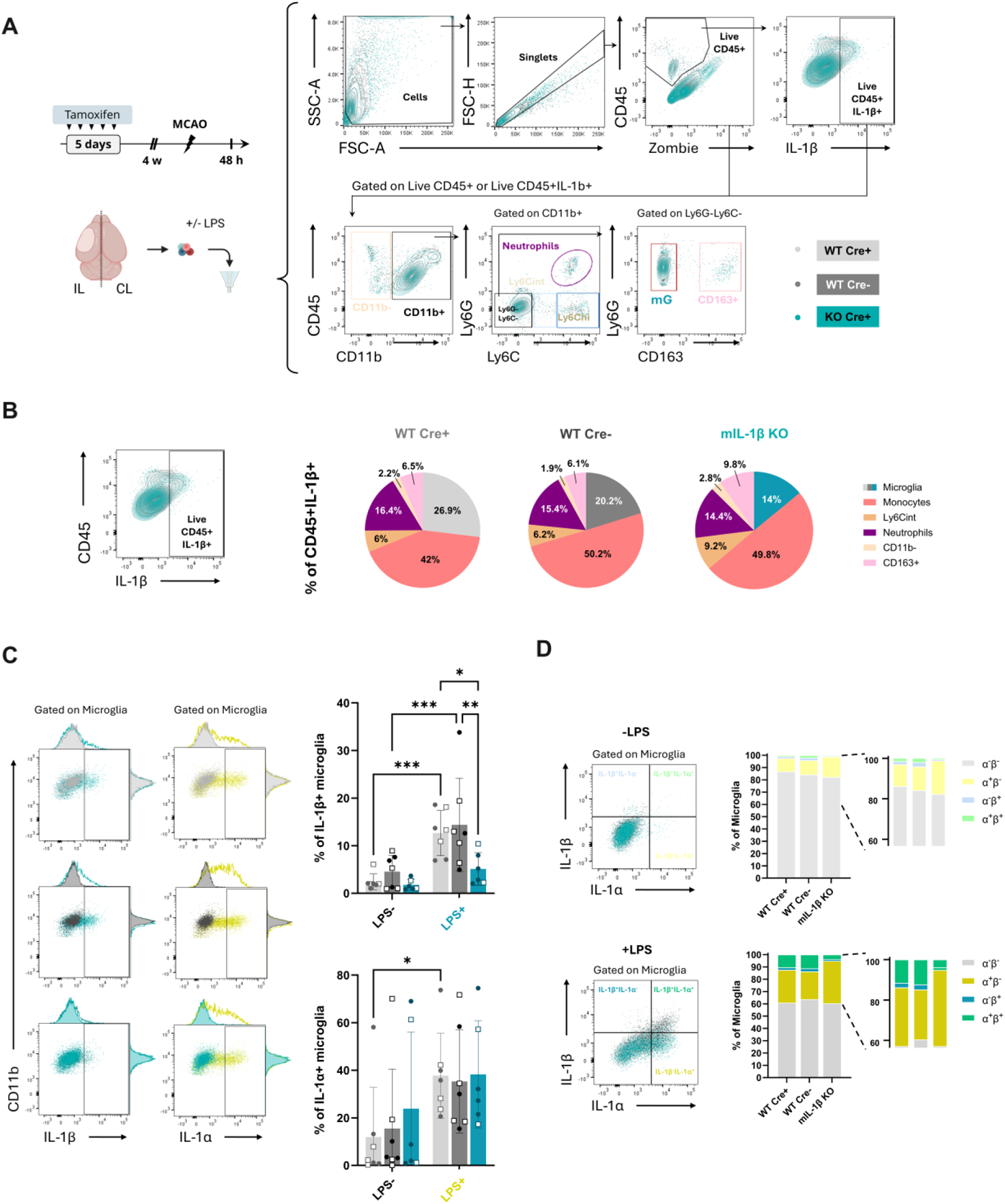
Microglial IL-1β deletion confirmation and effect on overall IL-1β expression and microglial IL-1α at 24 h post-MCAO. **A)** Experimental design, brain samples processing for flow cytometry and gating strategy. B) Representative gate of IL-1β+ cells within the Live CD45+ cell population (left) and pie charts (left) showing the distribution of IL-1β+ immune cells, shown as the mean percentage of cell populations in ipsilateral brain samples across genotypes. C) Representative IL-1β (left) and IL-1α (right) gating on microglia with or without LPS stimulation across genotypes, and graphs representing the proportion of IL-1β+ (top) and IL-1α+ (bottom) microglia within the microglial population in ipsilateral brain samples across genotypes (n=6-7/group, 2-way ANOVA followed by Šídák’s multiple comparisons test, *p<0.05, **p<0.01, ***p<0.001). D) Gating (left) and graphs (right) representing the proportion of IL-1α-IL-1β-, IL-1α-IL-1β+, IL-1α+IL-1β- and IL-1α+IL-1β+ cells within the microglial population in ipsilateral brain samples 24 h post-MCAO without (top) or with (bottom) LPS stimulation.

### 2.2. Focal cerebral ischemia

Transient focal cerebral ischemia was induced using the intraluminal model of proximal MCAO, as previously described^12^. Briefly, mice were anesthetized with isoflurane (4% for induction and 2% for maintenance in 30:70% mixture of N_2O_/O_2)_, a midline incision was made on the ventral surface of the neck and the left common carotid artery isolated and ligated. A 6-0 monofilament (Doccol, Sharon, MA, USA) was introduced into the external carotid artery. The filament was advanced approximately 10 mm into the common carotid with the filament making its way distal to the carotid bifurcation, to occlude the MCA and where it was left in place for 45 min. A 10 mm mark was made on the filament to visualize the required length to be inserted beyond the carotid bifurcation.

Body temperature was monitored and maintained at 37±0.5 °C throughout both procedures using a homoeothermic blanket and rectal probe (Harvard Apparatus, UK). Topical anesthetic (EMLA, 5% prilocaine and lidocaine, AstraZeneca, UK) was applied to skin incision sites prior to incision. Buprenorphine (0.05 mg/kg) was administered subcutaneously at the time of surgery. Mice were weighed daily for the first week post-stroke, given mashed diet and administered with 0.5 mL saline daily until body weight stabilized (approximately at 4 days post-stroke).

### 2.3. Functional tests

Neurological focal deficits were assessed at 48 h, 7 and 14 days post-stroke using a 28-point neurological scoring system previously described^13^. This is a cumulative score based on the following categories: body symmetry, gait, climbing, circling, front limb symmetry, compulsory circling, and whisker response. Animals are ranked from 0 (normal) to 4 (extreme deficit) for each category.

The affected (contralateral side to the stroke) forelimb grip strength was assessed as previously described^14^. Habituation to the test was performed 3 days prior to surgery to avoid neophobic behaviors, and grip strength was then recorded as the mean of at least 3 repeated measures at baseline and at 48 h, 7 and 14 days after MCAO.

### 2.4. Bromodeoxyuridine administration

Bromodeoxyuridine (BrdU) (Sigma-Aldrich) was dissolved in 0.9% NaCl and administered intraperitoneally with a single daily dose of 50 mg/kg from day 5 to day 14 post-MCAO, to track cell proliferation avoiding prominent immune cell infiltration within the first 5 days post-MCAO.

### 2.5. Tissue processing

For flow cytometry, anesthetized mice were transcardially perfused at 24 h post-MCAO with cold sterile PBS 0.1 M (pH 7.4). Brains were removed and ipsilateral and contralateral hemispheres kept in cold HBSS until processed.

For histological staining and immunostaining, anesthetized mice were transcardially perfused at 48 h or 14 days post-MCAO with cold saline followed by 50 mL of 4% paraformaldehyde (PBS 0.1 M, pH 7.4). Brains were removed, post-fixed in 4% paraformaldehyde and cryoprotected in sucrose 30% for 48-72 h, and snap-frozen in isopentane at -40°C. Brains were then cut to a thickness of 30 μm using a freezing sledge microtome (Bright Instruments, UK) and then stored in cryoprotectant (0.05 M Na_2H_PO_4_*2H_2O_, 5 mM NaH_2P_O_4,_ 30% anhydrous ethylene glycol, 20% glycerol) at -20°C until required.

### 2.6. Cresyl Violet stain

Cresyl violet staining was performed to measure lesion volume after MCAO, as previously described^15^. Briefly, 30 μm-thick brain sections were stained with 1% cresyl violet, and cover-slipped with DPX mounting medium (06522, Sigma-Aldrich). For each brain, infarct volumes were measured on defined coronal sections across the whole brain, spaced at approximately 360 µm apart (using image J). Coronal sections with their brain co-ordinates and lesion were integrated to estimate total lesion volume for each brain and corrected for edema.

### 2.7. IgG staining

To assess blood-brain barrier (BBB) permeability, IgG in the brain was stained by peroxidase-based immunohistochemistry. Endogenous peroxidase activity was quenched in 0.3% H_2_0_2_ and non-specific staining was blocked in Dulbecco’s phosphate buffeted saline (DPBS) with 2% donkey serum, 1% BSA and 0.3% Triton-X-100, and sections incubated in biotinylated anti-mouse IgG secondary antibody (1:500, A90-137B, Cambridge Bioscience) for 90 min at room temperature. Sections were then washed and incubated with avidin-biotin complex (ABC, PK-6100, Vector Laboratories) and color-developed using a diaminobenzidine solution (DAB, SK-4100, Vector Laboratories), following manufacturer’s instructions. To ensure comparability of DAB staining, all reactions were performed at the same time, with the same batch of DAB, with equal thickness sections exposed to DAB for the same amount of time. IgG staining intensity (using ImageJ) was averaged over coronal sections across the whole brain (at the same levels as for cresyl violet staining), with intensity increases in the ipsilateral hemisphere expressed as percentage increase compared to equivalent areas in the contralateral hemisphere. All sections were imaged at the same time with the same settings, and with no adjustment to brightness or contrast.

### 2.8. TUNEL assay

Apoptotic cells were detected using the TUNEL assay, following the kit instructions (Click-iT™ Plus TUNEL Assay, C10617 Alexa Fluor™ 488, Invitrogen™). This was done on the previously described 30 μm thick coronal brain sections, which for the purpose of this protocol were mounted onto the slide beforehand. The TUNEL assay was performed in combination with the primary antibody anti-Ionized calcium-binding adaptor molecule 1 (Iba1) (1:500, ab178846, Abcam), and DAPI (25 ng/mL in dH₂O) was used to detect cellular nuclei. TUNEL-positive cells were quantified, and co-localization analysis was performed with Iba1+ cells (Alexa Fluor™ 647 Secondary Antibody, A31573, ThermoFisher Scientific, 1:500).

### 2.9. Immunofluorescence

Brain sections were washed in DPBS with 0.1% Tween20. For BrdU immunostaining, brain sections were then incubated in 2M HCl in DPBS for 1 h to facilitate antibody access to the DNA-incorporated BrdU, followed by neutralization with 0.1M borate buffer for 10 min. All sections were incubated in blocking solution (1% BSA and 5% donkey serum in DPBS with 0.05% Tween20, 0.1% Triton X-100 and 0.2M glycine) for 1 h and incubated overnight at 4°C with the following primary antibodies (in 1% BSA and 0.3% Triton X-100 in PBS): rat-anti-BrdU (1:100, ab6326, Abcam), goat-anti-CD13 (1:50, AF2335, R&D Systems), rat-anti-CD31 (1:200, 553370, BD Biosciences), rabbit-anti-doublecortin (DCX) (1:200, ab18723, Abcam), rat–anti-Lymphocyte antigen 6 complex locus G6D (Ly6G) (1:750, 127602, Biolegend), anti-Iba1 (1:1000, ab178846, Abcam), and chicken-anti-glial fibrillary acidic protein (GFAP) (1:1000, A85307, Antibodies.com), mouse-anti-NeuN (1:1000, A85405, Antibodies.com). The following secondary antibodies were incubated for 2 h at room temperature: Alexa Fluor 488 donkey anti-rat (10 µg/mL; A21208), Alexa Fluor 488 donkey anti-goat (10 µg/mL; A32814), Alexa Fluor 555 donkey anti-rabbit (10 µg/mL; A31572), Alexa Fluor 647 donkey anti-mouse (10 µg/mL; A31571), and Alexa Fluor 647 donkey anti-rat (10 µg/mL; A48272). Sections were counterstained with DAPI (25 ng/mL in dH₂O) to stain for cell nuclei and mounted with Prolong Gold antifade mounting medium (Thermo Fisher Scientific, P36934).

Images were collected on a 3D-Histech Pannoramic-250 microscope slide-scanner using a x20/0.30 Plan Achromat objective (Zeiss, Germany). All quantitative analyses were performed on at least 2 regions of interest (ROIs) per defined brain area, in 3 different coronal brain slices per animal, at equivalent coordinates relative to Bregma.

### 2.10. Image analysis

Images were captured using a Zeiss Axiomager.M2 upright microscope, a 20X Plan Apochromat objective and a CoolSnap HQ2 camera (Photometrics). This was operated via Micro-Manager software (v1.4.23). Analyses were performed in at least 3 coronal brain sections per animal, between bregma -2.2 mm to +0.52 mm. ROIs were determined based on DAPI counterstain and anatomical landmarks. Peri-infarct ROIs were drawn in the ipsilateral hemisphere (IL) and reflected to the contralateral hemisphere (CL).

For the analysis of angiogenesis and pericyte coverage (CD31+, CD13+), 2 square ROIs were taken in the peri-infarct cortex and 3-4 square ROIs in the peri-infarct sub-cortex, and reflected to the contralateral hemisphere, as shown in Figures 5 and S4. Exported images of stained vessels (CD31+) were analyzed using AngioTool64 (Version 0.6a, 0.2.18.14), an open-source software for quantitative angiogenesis analysis (CD31 images). After outlining the vessel network, AngioTool computes several morphological and spatial parameters including (i) the overall size of the vascular network, (ii) total and average vessel length, (iii) vascular density, (iv) lacunarity, indicative of vessel non-uniformity, and (v) branching index (branch points / unit area), indicative of angiogenic sprouting activity. Image J was used for the analysis of pericyte density (CD13+ area), and coverage (%CD13+ area/CD31+ area), on CD13+ images, as well as for the analysis of neutrophil density (Ly6G+) and microglia (Iba1+).

QuPath (v0.5.1) was used for the analysis of TUNEL staining and neurogenesis markers. All parameters were optimized from representative samples and applied uniformly, with thresholds constant across experimental groups. Classifiers were trained using at least 3 training images and with at least 100 detections per classification, using the Qupath Random Trees Model. For the TUNEL analysis, cell detection was based on TUNEL+ cells, followed by co-colocalization with Iba1+ signal. For the neurogenesis markers analysis, nuclear cell detection was based on DAPI and NeuN+, followed by co-localization analyses with DCX+ and BrdU+.

### 2.11. Flow cytometry

Brain single-cell suspensions were stimulated with 50 ng/ml bacterial lipopolysaccharides (LPS) (InvivoGen; tlrl-3pelps) in the presence of Brefeldin A (Biolegend) at 37°C or kept in RPMI 10% FBS at 4°C. After 3 h, cells were washed in PBS and incubated with antibody cocktails plus anti-CD16/32 (2.4G2; BioXcell) for 15-20 min at 4°C. Cells were stained with combinations of the following antibodies, which were obtained from eBioscience and BioLegend: CD11b (M1/70), TCR-b (H57-597), MHC-II (M5/114.15.2), Ly6G (1A8), CD45 (30-F11), B220 (RA3-6B2), Ly6C (HK1.4), CD163 (S15049I), IL-1α (ALF-161) and IL-1β (NJTEN3). Dead cells were excluded by use of a Live/Dead Fixable blue dead cell stain kit (Molecular Probes). Following surface staining, cells were fixed in using the eBioscience Foxp3/Transcription Factor Stainign Buffer Set as per the manufacturer’s instructions and washed in eBioscience Foxp3 Permeabilization Buffer. For intracellular staining, samples were stained overnight at 4°C in the dark with fluorochrome-conjugated antibodies in eBioscience Foxp3 Permeabilization Buffer. After washing and resuspending in PBS, samples were acquired using a LSRFortessa (BD Biosciences) flow cytometer and analyzed with FlowJo (Treestar).

### 2.12. LEGENDplex assay

Prior transcardiac perfusion for histological staining at 24 h, 48 h or 14 days post-MCAO, venous blood was obtained by cardiac puncture from the right ventricle, into 3.8% sodium citrate pre-loaded syringes (2% of the final volume). Blood samples were then centrifuged at 1500 rcf for 15 min to obtain plasma, which was stored at -70°C until used.

Plasma cytokines were simultaneously quantified using a mouse inflammation LEGENDplex 13-plex custom panel (Biolegend, 740446), including IL-1α, IL-1β, IL-6, IL-10, IL-12p70, IL-17A, IL-23, IL-27, monocyte chemoattractant protein-1 (MCP-1), interferon β (IFN-β), interferon gamma (IFN-γ), tumor necrosis factor α (TNF-α), and granulocyte-macrophage colony-stimulating factor (GM-CSF). The assay was performed according to the manufacturer’s instructions in V-bottom plates and analyzed using a BD Symphony flow cytometer (BD Biosciences). The data were analyzed using Qognit LEGENDplex (Biolegend) data analysis software.

### 2.13. Statistical analysis

Statistical analyses were performed using GraphPad Prism 8.0. All values are expressed as mean ± standard deviation (SD) or median (InterQuartile Range, IQR) according to the normal or non-normal distribution of the represented variable, respectively. The normality of continuous variables was assessed using the Shapiro-Wilk test (n < 30) or Kolmogorov-Smirnov test (n ≥ 30). Data were analyzed using unpaired t-test, one-way ANOVA (followed by Dunnet’s multiple comparison post hoc test), two-way ANOVA (followed by Sidak’s multiple comparison post hoc test), or by fitting a mixed effects model using Restricted Maximum Likelihood (REML), followed by Sidak’s multiple comparison post hoc test. The Kruskal Wallis test (followed by Dunn’s multiple comparisons post hoc test) was used for non-normally distributed variables. Survival analyses were performed using the Gehan-Breslow-Wilcoxon test. The significant level was set at p<0.05, and statistical trend reported for values 0.05≤p<0.1.

## 3. Results

### 3.1. Deletion of microglial IL-1β does not affect IL-1β expression in other immune cell sources nor microglial IL-1α expression at 24 h after stroke

First, we investigated the effects of microglial IL-1β deletion on the expression of IL-1α and IL-1β in the post-stroke brain (Figure 1A). To characterize the cellular source of IL-1β, we analyzed the proportion of live CD45^+^IL-1β^+^ cell populations across genotypes (Figure 1B). We found that the immune cell populations expressing IL-1β in the post-stroke brain were comparable across genotypes, and were predominantly comprised of infiltrating monocytes, followed by microglia and neutrophils. The remaining source of CD45+IL-1β+ cells were CD11b+Ly6G-Ly6Cint, CD163+ border-associated macrophages (BAMs, CD11b+Ly6G-Ly6C-CD163+) and a small proportion of CD11b-cells. In contrast, microglia accounted for most of the cellular source of IL-1α in the post-stroke brain across all genotypes, followed by infiltrating monocytes (Figure S1A).

We then analyzed the proportion of IL-1β+ cells within the microglial population (Figure 1C), showing that only up to 5% of microglia were IL-1β+ in the ipsilateral brain at 24 h post-MCAO, with no significant differences between groups. Upon LPS stimulation, the percentage of IL-1β+ microglia was significantly increased by over 10% in both WT Cre+ (p<0.001 vs. LPS-) and WT Cre-(p<0.001 vs. LPS-), while the expression of IL-1β+ in the mIL-1β KO group remained unchanged (p>0.05 vs. LPS-). Furthermore, the proportion of IL-1β+ microglia in LPS stimulated ipsilateral brain samples, collected at 24 h post-MCAO, was significantly lower in the mIL-1β KO group, compared to both the WT Cre+ and WT Cre-groups (p<0.05 and p<0.01, respectively). These results suggest a partial yet successful Cre recombinase-mediated deletion of IL-1β in microglial cells at 24 h after MCAO. Importantly, there were no significant differences in neither LPS-unstimulated or LPS-stimulated ipsilateral brain samples between WT Cre+ and WT Cre-groups.

Notably, successful deletion of microglial IL-1β did not affect the expression of microglial IL-1α, as seen by the proportion of IL-1α+ microglia in the same post-stroke brain samples with or without LPS priming. Furthermoe, upon LPS stimulation, the proportion of IL-1α+ microglia was only modestly increased (p=0.0424, p=0.1150 and p=0.1051 in WT Cre+, WT Cre- and mIL-1β KO, respectively, vs. LPS-group) yet to comparable levels between groups (Figure 1C).

These results confirm earlier observations in different ischemic stroke models showing that IL-1α is rapidly expressed in the brain after ischemic stroke exclusively in microglia, followed by the expression of IL-1β in a small subset of microglia, but also in infiltrating immune cells from the periphery, such as neutrophils. We then asked whether IL-1β-expressing microglia also express IL-1α and vice versa, or whether post-stroke microglia specialize to exclusively express either IL-1β or IL-1α (Figure 1D). We found that over 80% of all microglia in the ipsilateral brain 24 h post-MCAO were negative for both IL-1α and IL-1β, over 10% of microglia expressed IL-1α but not IL-1β, while fewer microglia were double positive for IL-1α and IL-1β or only expressed IL-1β.

### 3.2. Microglial IL-1β deletion does not affect brain immune cell recruitment nor microglial activation at 24 h after stroke

Next, we investigated the effects of microglial IL-1β deletion on the acute immune cell profile in the post-stroke brain. The proportion of immune cell populations (CD45+) at 24 h after MCAO remained unchanged across genotypes (Figure S1B). Most immune cell populations in the post-stroke brain were microglia (CD45+CD11b+Ly6G-Ly6C-CD163-), followed by infiltrating monocytes (CD45+CD11b+Ly6G-Ly6Chi), and neutrophils (CD45+CD11b+Ly6G+Ly6C-). Interestingly, when splitting the analyses by sex, the proportion of infiltrating neutrophils was reduced in female mIL-1β KO mice (p<0.05 vs. WT Cre-) but not in male mIL-1β KO mice (Figure S1C).

To assess potential differences in microglial activation, reflected by the expression of CD45, CD11b or MHC II, we analyzed the mean fluorescence intensity (MFI) of such markers, revealing no significant differences across genotypes (Figure S1D). Importantly, no differences in immune cell frequencies nor microglial MHC II, CD45 nor CD11b expression were observed between WT Cre+ and WT Cre-mice.

### 3.3. Microglial IL-1β contributes to acute stroke outcome and neutrophil recruitment

Having discarded a potential monoallelic effect of CX3CR1 on immune responses in our initial immune profiling, and proving no differences in neurological outcome nor infarct size between WT Cre+ and WT Cre-mice (Figure S2), all further studies were performed using a tamoxifen-treated WT Cre-, as previously reported^6^.

At 48 h post-stroke, microglial IL-1β deletion resulted in improved neurological function, as seen by a significant reduction in the 28-point neuroscore (p<0.05 vs. WT Cre-; Figure 2B), alongside a modest non-significant, reduction in infarct size (p<0.10 vs. WT Cre-; Figure 2C), and BBB breakdown (p<0.10 vs. WT Cre-; Figure 2D). Notably, when splitting the analyses by sex, improved neurological outcome was observed in female (p<0.05 vs. WT Cre-) but not male mIL-1β KO mice at 48 h post-MCAO. Similarly, infarct size was significantly reduced in female (p<0.05 vs. WT Cre-) but not male mIL-1β KO mice. Quantification of circulating cytokines at 48 h post-stroke revealed reduced levels of IL-1α (p<0.01), TNFα (p<0.05), IL-10 (p<0.01), and IL-6 (p<0.01) upon microglial IL-1β deletion vs. WT Cre- (Table 1), and a moderate non-significant reduction of MCP-1 (p<0.10) and IL-27 (p<0.10) vs. WT Cre-, (Table 1). Interestingly, when splitting the analyses by sex, only females showed a significant reduction in IL-10 and IL-6 upon microglial IL-1β deletion (p<0.05 vs. WT Cre-, respectively), while surprisingly, IL-17A levels were significantly lower in the mIL-1β KO female mice (p<0.01 vs. WT Cre-), but higher in mIL-1β KO male mice (p<0.10 vs. WT Cre-); Figure S3B. No differences in circulating cytokine levels were observed between groups at 24 h nor 14 days post-MCAO (Tables S1, S2).

**Figure 2.**
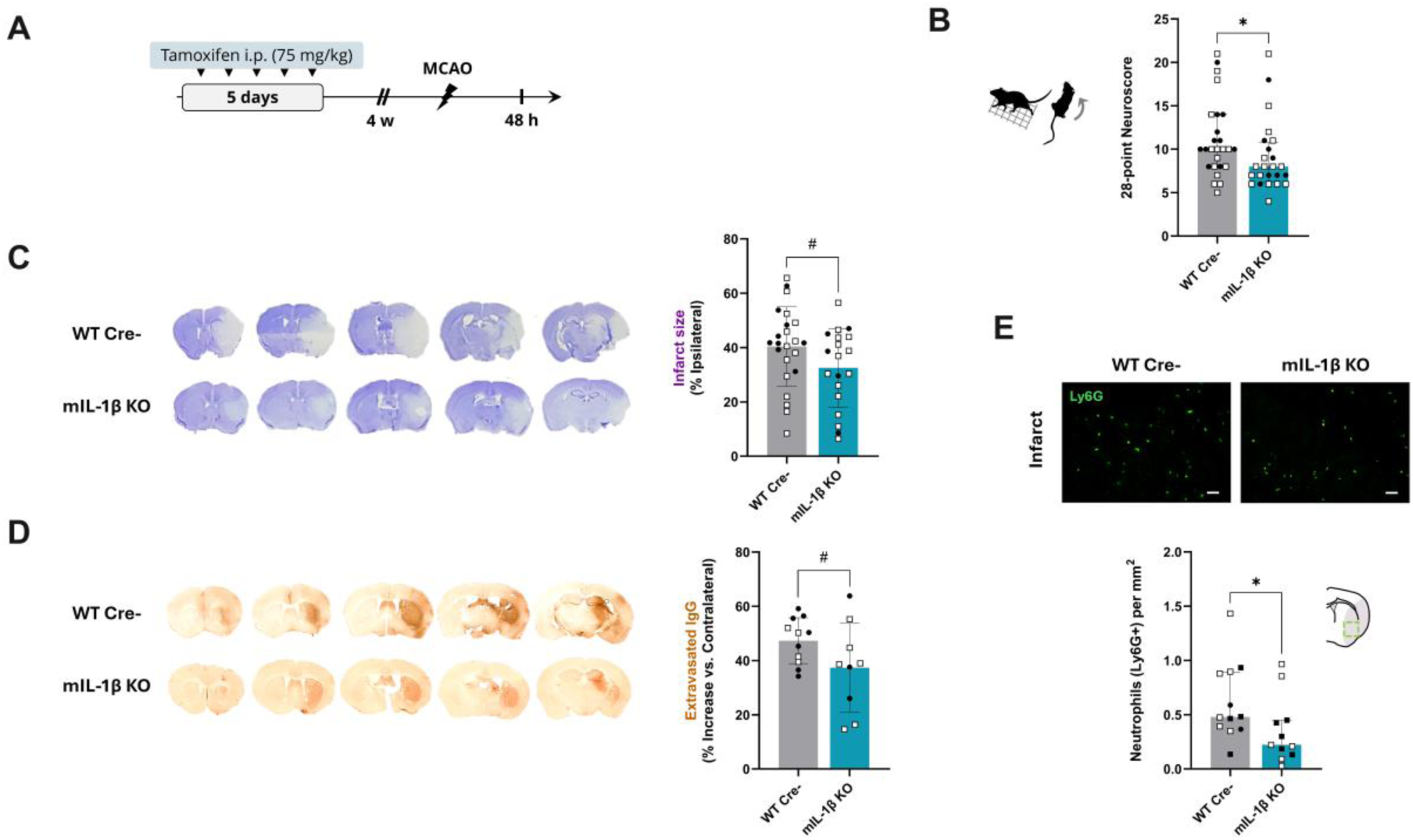
Microglial IL-1β deletion improves outcome and reduces neutrophil recruitment at 48 h post-stroke. A) Schematic representation of the experimental design. B) Neurological outcome at 48 h post-MCAO assessed by a 28-point based neurological score (n=23-24, Mann Whitney test). C) Representative cresyl violet-stained brains and infarct size quantification expressed as the percentage of total ipsilateral hemisphere volume (n=20-23, unpaired t-test). D) Representative IgG IHC-stained brains and IgG parenchymal extravasation quantification, expressed as the percentage of intensity increase in the ipsilateral versus the contralateral hemisphere (n=9-11, unpaired t-test). E) Representative immunostaining of the infiltrating neutrophils (Ly6G+) in the infarct region (scale bar: 10 µm, schematic of the ipsilateral brain hemisphere depicting the infarct region in grey and representative ROI) and quantification of neutrophil density (n=11-12/group, Mann Whitney test). #p<0.10, *p<0.05.

**Table 1.** Plasma cytokine levels at 48 h after MCAO.

| Cytokines | WT<br>(pg/mL) | mIL-1β KO<br>(pg/mL) | t-test/M-W<br>(p-value) |
| --- | --- | --- | --- |
| IL-23 | 108.6 (1795) | 112.2 (147.2) | 0.6461 |
| IL-1α | 1.68±0.67 | 1.13±0.41 | **0.0070** |
| TNFα | 9.18 (17.16) | 5.49 (20.15) | **0.0219** |
| IFNγ | 1.90 (5.0) | 2.01 (5.8) | 0.7078 |
| MCP-1 | 14.73 (39.1) | 10.0 (18.1) | 0.0630 |
| IL-12p70 | 2.3 (7.9) | 3.3 (15.5) | 0.6375 |
| IL-1β | 7.9 (29.4) | 7.9 (5.6) | 0.6027 |
| IL-10 | 103.2±59 | 51.2±30.6 | **0.0088** |
| IL-6 | 7.62 (23.4) | 2.0 (6.4) | **0.0013** |
| IL-27 | 81.6 (466) | 49.6 (91.6) | 0.0532 |
| IL-17A | 1.63 (3.7) | 1.53 (4.7) | 0.8911 |
| IFN-β | 177.6 (192.3) | 112.1 (146.3) | 0.5556 |
| GM-CSF | 6.31 (8.3) | 6.94 (2.6) | 0.7214 |

Data represented as mean ± SD or median (IQR) for parametric or non-parametric analyses, respectively.

IL-1 and microglia are known to drive brain neutrophil recruitment after stroke, thereby limiting BBB disruption and secondary tissue injury.^16–18^ Based on this and given our observation that microglial IL-1β deletion improves neurological outcome with modest infarct reduction and BBB protection, we next assessed neutrophil accumulation in the ischemic brain. Our results showed significantly reduced neutrophil numbers within the infarct region of mIL-1β KO mice at 48 h post-MCAO (p<0.05 vs. WT Cre-; Figure 2E). We then determined whether changes in acute functional outcome, infarct size, systemic inflammation and neutrophil recruitment after stroke were accompanied by altered cell death within the infarcted area using TUNEL staining (Figure S3C). The density of TUNEL⁺ cells did not differ between genotypes in either cortical or subcortical infarct regions. Additionally, microglial cell death within the ischemic core, which has been previously linked to neutrophil accumulation in the post-stroke parenchyma,^18^ remained unchanged, as indicated by comparable Iba1+TUNEL+ cell numbers in both the cortical and subcortical infarct regions across genotypes (Figure S3D). The area covered by microglia (Iba1+) also remained unchanged across brain regions and between genotypes (Figure S3E).

### 3.4. Long-term post-stroke recovery is not affected by microglial IL-1β

To study the effect of microglial IL-1β on post-stroke recovery, WT Cre- and mIL-1β KO mice were followed up to 14 days post-MCAO (Figure 3A).

**Figure 3.**
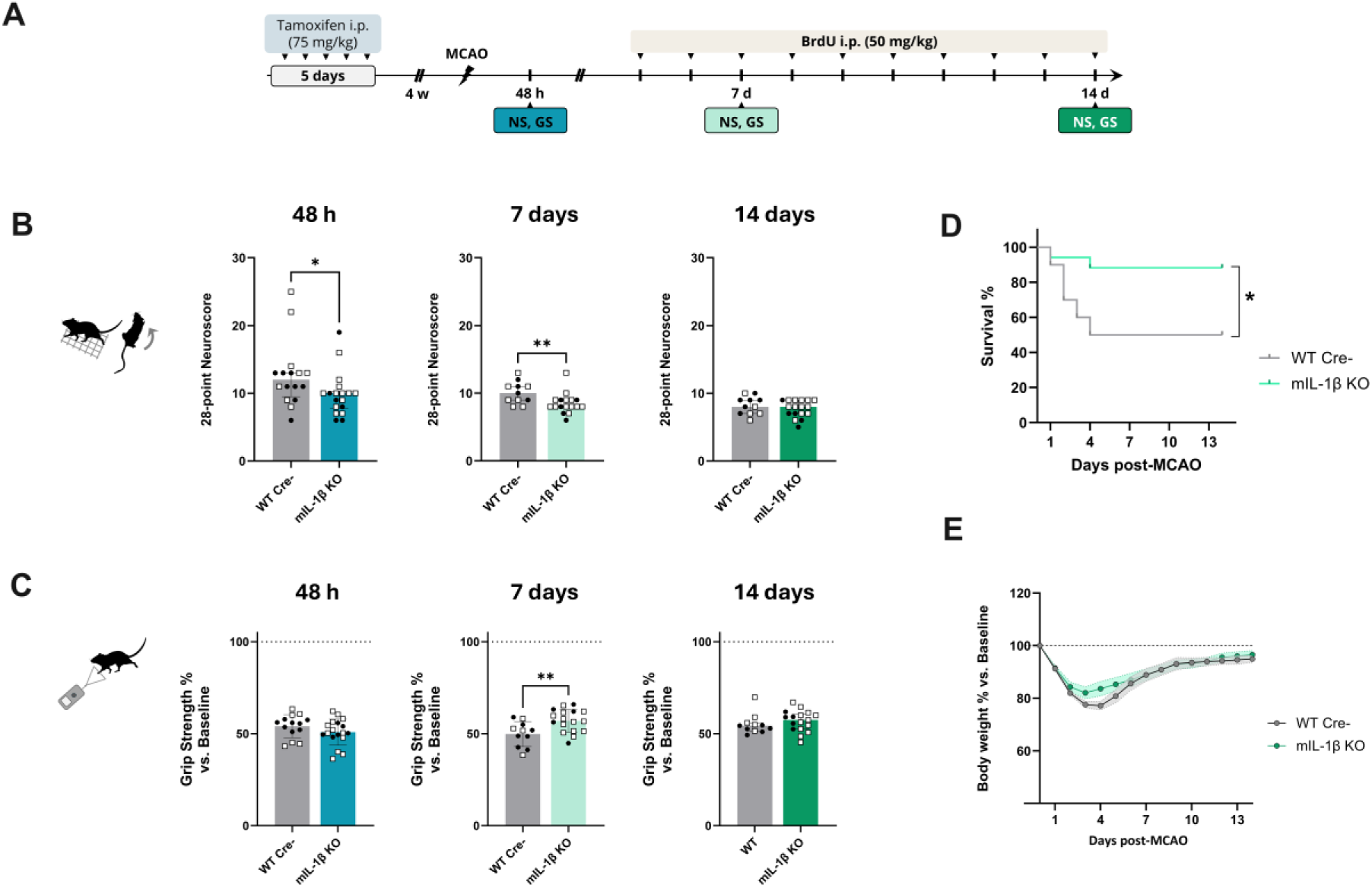
Microglial IL-1β deletion improves outcome up to 7 days after stroke. A) Schematic representation of the experimental design. B) Kaplan-meier curve showing survival proportions over time across groups (n=10-17 per group, Gehan-Breslow-Wilcoxon test). C) Neurological outcome at 2, 7 and 14 days post-MCAO assessed by a 28-point based neurological score (NS, n=16-18, 11-17, 11-17 per group, Mann Whitney test). D) Grip strength (GS) at 2, 7 and 14 days post-MCAO represented as the percentage of the corresponding baseline strength of the contralateral affected (left) front paw (n=16-18, 11-17, 11-17 per group, unpaired t test for comparisons at 2 and 7 days post-MCAO, Mann Whitney test for comparison at 14 days post-MCAO). E) Body weight expressed as the percentage of corresponding baseline over time across groups (n=18-19/group, mixed effects model using REML).

Again, microglial IL-1β deletion led to an improvement in focal neurological deficits, as seen by a reduction of the neurological score at 48 h (p<0.05 vs. WT Cre-), confirming the role of microglial IL-1β on acute stroke outcome. This neurological improvement was maintained at 7 days post-MCAO (p<0.01 vs. WT Cre-), however both groups recovered to similar neurological scores at 14 days after stroke (Figure 3B). Both WT Cre- and mIL-1β KO showed a substantial yet similar reduction of grip strength in the contralateral (left) affected front paw at 48 h post-MCAO (54.1% and 50.9% of baseline in WT Cre- and mIL-1β KO, respectively). However, in line with the neurological score results, mIL-1β KO mice showed an improved grip strength at 7 days post-stroke (57% and 49% vs. baseline in mIL-1β KO and WT Cre-, respectively; p<0.01), which was then comparable to that of the WT Cre-group by day 14 (Figure 3C).

The overall survival was significantly higher in the mIL-1β KO group (p<0.05 vs. WT Cre-). Attrition due to humane endpoint criteria occurred within the first 4 days post-stroke, leading to 50% and 88% survival at 14 days post-MCAO in the WT Cre- and mIL-1β KO groups, respectively (Figure 3D). Finally, alongside improved survival and acute functional outcome, mIL-1β KO mice seemed to start recovering from post-stroke body weight loss earlier than the WT Cre-group (Figure 3E). However, while there was a significant effect of body weight changes over time (p<0.0001), there were no differences between genotypes nor an interaction between time and genotype.

Altogether, our results suggest that microglial IL-1β may play a key role on acute stroke outcome but may not influence long-term post-stroke recovery.

### 3.5. Microglial IL-1β deletion differentially regulates subventricular zone and hippocampal neurogenesis

We next explored whether microglial IL-1β deletion affected post-stroke neurogenesis (Figure 4A, F). First, we investigated whether microglial-derived IL-1β affects the neurogenic pool in the subventricular zone (SVZ), ectopic post-stroke neuroblast migration, or the integration of newborn neurons within the peri-infarct territory. For this, we assessed the effect of microglial IL-1β deletion on NSC proliferation in the SVZ, as well as neuroblast migration and neuronal differentiation and incorporation in the peri-lesional regions at 14 days post-MCAO (Figure 4B, C). Neuroblast cell density (DCX+ cells) in the ipsilateral SVZ, normalized by the corresponding contralateral SVZ, indicated an increase in response to stroke and, interestingly, this increase was significantly higher in the mIL-1β KO group compared to the WT Cre-group (p<0.001). The proliferation of neuroblasts in the ipsilateral SVZ, assessed by the density of DCX+BrdU+ cells normalized to the contralateral SVZ, was also increased, although to the same extent in both the mIL-1β KO and WT Cre-groups (Figure 4D). To investigate the ectopic migration of SVZ-derived neuroblasts towards the stroke lesion, we quantified the density of DCX+ and DCX+BrdU+ cells in the peri-lesional subcortical region adjacent to the SVZ. The normalized ipsilateral DCX+ cell density was higher, although not reaching statistical significance, in the mIL-1β KO group compared to the WT Cre-group (p<0.10), while the ipsilateral DCX+BrdU+ cell density was significantly higher in the mIL-1β KO group than in the WT Cre-group (p<0.05; Figure 4E).

**Figure 4.**
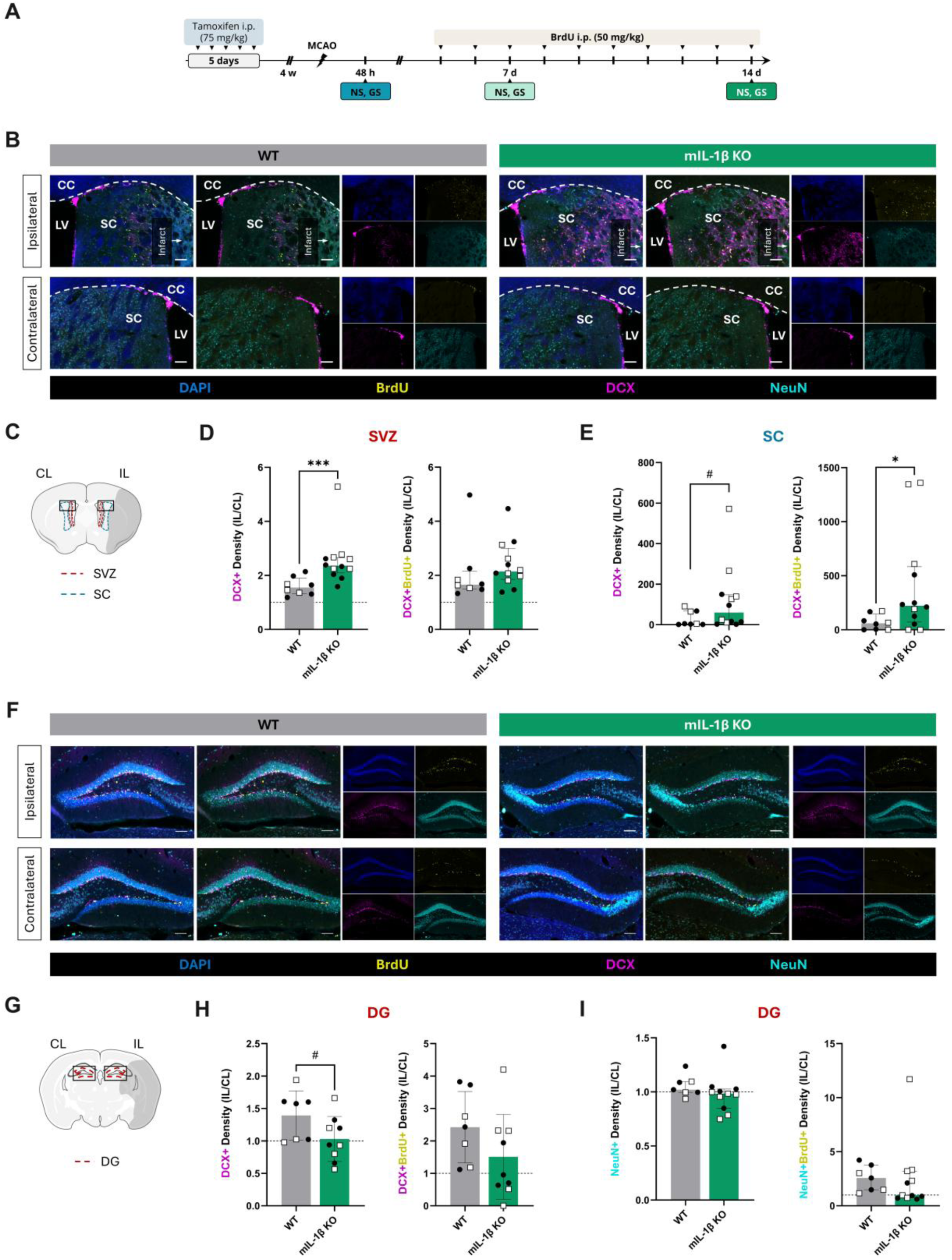
Microglial IL-1β deletion leads to opposite effects in different neurogenic niches. A) Schematic representation of the experimental design. B) Representative immunostaining of the neurogenic pool in the subventricular zone (SVZ) and ectopic migration towards the subcortical (SC) infarct region, stained for neuroblasts (DCX+), proliferating cells (BrdU+) and mature neurons (NeuN+), in the ipsilateral (IL) and contralateral (CL) hemispheres of WT and mIL-1β KO mice (scale bar: 100 µm). CC: corpus callosum, LV: lateral ventricle. C) Schematic of a coronal brain section at bregma +0.3 showing ROIs selected for analyses of the SVZ and SC areas, depicting the infarct region in grey and location of the representative immunofluorescence microscopy images in (B). D) Quantification of DCX+ cell density (left) and DCX+BrdU+ cell density (right) in the IL SVZ normalized by the respective CL ROI (n=8-12/group, Mann Whitney test, ***p<0.001). E) Quantification of DCX+ cell density (left) and DCX+BrdU+ cell density (right) in the IL SC normalized by the respective contralateral ROI (n=8-12/group, Mann Whitney test, #p<0.10, *p<0.05). F) Representative immunostaining of the hippocampal neurogenic pool in the DG, stained for neuroblasts (DCX+), proliferating cells (BrdU+) and mature neurons (NeuN+), in the IL and CL hemispheres of WT and mIL-1β KO mice (scale bar: 100 µm). G) Schematic of a coronal brain section at bregma -2.0 showing ROIs selected for analyses of the DG, depicting the infarct region in grey and location of the representative immunofluorescence microscopy images in (F). H) Quantification of DCX+ cell density (left) and DCX+BrdU+ cell density (right) in the IL DG normalized by the respective CL ROI (n=7-9/group, unpaired t-test, p<0.10). I) Quantification of NeuN+ cell density (left) and NeuN+BrdU+ cell density (right) in the IL DG normalized by the respective contralateral ROI (n=7-9/group, Mann Whitney test).

Next, we investigated whether microglial-derived IL-1β affects endogenous hippocampal neurogenesis by assessing the neurogenic pool in the subgranular zone (SGZ) and the integration of newborn neurons in the granular cell layer (GCL) of the dentate gyrus at 14 days post-MCAO (Figure 4F, G). As expected, the DCX+ cell density was increased in the ipsilateral SGZ in the WT Cre-group, which was not seen with microglial IL-1β deletion (p<0.10 vs. WT Cre-), while there were no differences in the density of DCX+BrdU+ cells normalized to the contralateral SGZ (Figure 4H). Both mIL-1β KO and WT Cre-mice showed a similar increase in the incorporation of newborn neurons, identified as NeuN+BrdU+ cells, in the ipsilateral GCL. However, the incorporation of newborn neurons did not affect overall neuronal density (NeuN+), which was similar between genotypes (Figure 4I).

Altogether, these results suggest that microglial IL-1β deletion increases the neurogenic pool in the subventricular zone and ectopic neuroblast migration in the subcortical peri-lesional area, while limiting post-stroke hippocampal neurogenesis at 14 days post-stroke.

### 3.6. Post-stroke angiogenesis and astrogliogenesis are not affected by microglial IL-1β deletion

We previously showed that conditional deletion of microglial IL-1α reduces peri-infarct angiogenesis and astrogliosis and impairs functional recovery 14 days after MCAO^6^. We therefore asked whether microglial IL-1β similarly contributes to these post-stroke neurorepair processes (Figure 5A, B). In line with our previous findings in this model^6^, subcortical, but not cortical (Figure S4), peri-infarct vascular density was enhanced in both WT Cre- (p<0.0001 vs. contralateral) and mIL-1β KO mice (p<0.001 vs. contralateral). The total vessels length was also enhanced in peri-infarct subcortical areas in the WT Cre- and mIL-1β KO groups (p<0.01 vs. contralateral, respectively). Analysis of junctions’ density revealed significant differences across genotypes, specifically a significant increase in the subcortical peri-infarct area of mIL-1β KO mice (p<0.05 vs. WT Cre-), yet no significant differences were identified in this region compared to its corresponding contralateral hemisphere in neither WT Cre-nor mIL-1β KO groups. The total number of vessel endpoints was unchanged across hemispheres and genotypes (p>0.05 for all comparisons). Finally, lacunarity was significantly lower in the peri-infarct subcortical area in both WT Cre- and mIL-1β KO groups (p<0.001 vs. contralateral, respectively) but was similar between groups (Table 2, Figure 5C, D). Pericytes have been involved in neovascularization, vessel stabilization and BBB integrity after cerebral ischemia^19^. Thus, alongside vessel network analyses, and considering the modest BBB damage seen acutely after stroke upon microglial IL-1β deletion, we also assessed pericyte density and coverage in the peri-infarct and corresponding contralateral areas. Both pericyte density and coverage were increased in the peri-infarct subcortical and cortical region in both WT Cre- and mIL-1β KO (p<0.0001 vs. contralateral, respectively), while no differences were observed between genotypes (Figure 5E, Figure S4E). Finally, microglial IL-1β depletion did not influence the density of reactive astrocytes, measured by the GFAP+ area in the subcortical peri-infarct area (Figure 5F). Altogether, our results confirm endogenous post-stroke angiogenesis and astrogliosis occurring at 14 days post-MCAO, however these do not seem to be influenced by microglial IL-1β.

**Figure 5.**
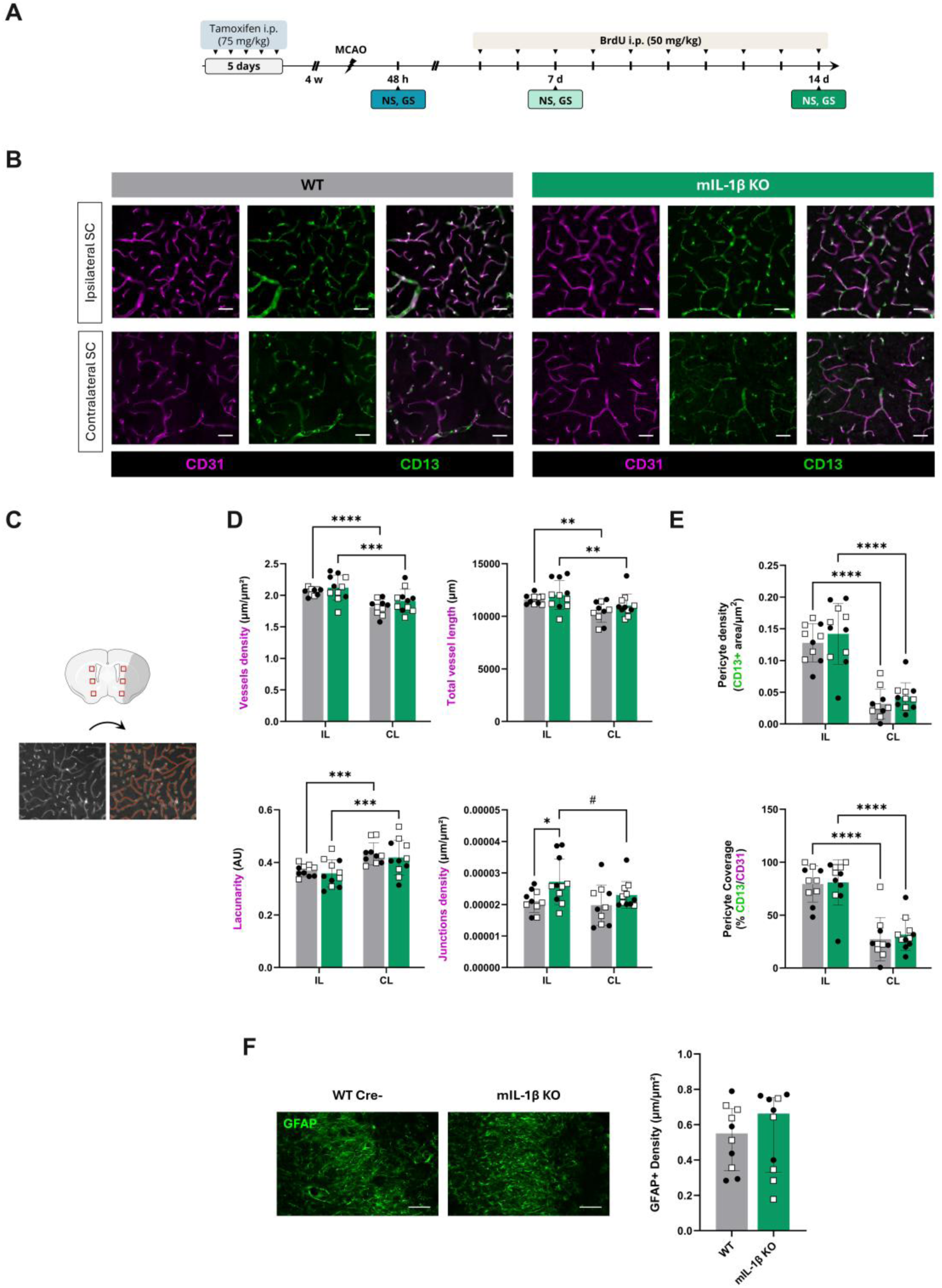
Microglial IL-1β deletion does not affect post-stroke angiogenesis or astrogliosis. A) Schematic representation of the experimental design. B) Representative immunostaining of the vasculature, stained for CD31 (endothelial cells) and CD13 (pericytes) in the peri-infarct subcortical region (SC) in the ipsilateral (IL) hemisphere and reflective contralateral (CL) SC region of WT and mIL-1β KO mice (scale bar: 50 µm). C) Schematic of a coronal brain section at bregma +0.3 showing ROIs selected for analyses of the SC areas, depicting the infarct region in grey (top) and representative CD31-stain image for vessel network analysis with AngioTool (left) and output image (right) showing the outline of the vasculature (yellow), skeleton representation of the vasculature (red) and branching points (blue). D) Graphs presenting selected vessel network analysis outputs, including vessels density, total vessel length, lacunarity and junctions density (n=10-11/group, 2-way ANOVA, and REML for junctions density analysis, followed by Šídák’s multiple comparisons test, #p<0.10, *p<0.05, **p<0.01, ***p<0.001, ****p<0.0001). E) Quantification of pericyte (CD13+) cell density (top) and pericyte coverage, expressed as the percentage of CD13+ area per CD31+ vessels area, in the IL and CL peri-infarct SC areas (n=8-12/group, 2-way ANOVA followed by Šídák’s multiple comparisons test, #p<0.10, *p<0.05). F) Representative immunostaining of the glial scar (GFAP+) in the IL SC region (left) and quantification of reactive astrocyte (GFAP+) density (scale bar: 100 µm, n=10/group, Mann Whitney test).

**Table 2.** Vessel network analysis at 14 days post-MCAO.

| Subcortical |  |  |  |  |  |  |
| --- | --- | --- | --- | --- | --- | --- |
|  | *Ipsilateral* |  | *Contralateral* |  | 2w ANOVA /<br>REML<br>(p-value) | Šídák's post-<br>hoc test<br>(p-value; IL/CL) |
|  | WT | mIL-1β KO | WT | mIL-1β KO |  |  |
| Junctions density<br>(μm/μm²) | 2.1E-05<br>(5.6E-06)<br>ns | 2.6E-05<br>(1.3E-05)<br># | 1.9 E-05<br>(1.0E-05) | 2.2E-05<br>(5.1E-06) | H: **0.0427**<br>G: **0.0378**<br>H x G: 0.2579 | **0.0323** / 0.3538 |
| Vessels density<br>(μm/μm²) | 2.05±0.06<br>**** | 2.12±0.20<br>*** | 1.81±0.12 | 1.91±0.18 | H: **<0.0001**<br>G: 0.1796<br>H x G: 0.5779 | 0.5421 / 0.3538 |
| Total length<br>(μm) | 11593±454<br>** | 12030±1370<br>** | 10412±998 | 11039±1101 | H: **<0.0001**<br>G: 0.2005<br>H x G: 0.6723 | 0.5723 / 0.3538 |
| Average vessels length<br>(μm) | 308±46<br>** | 313±62<br>ns | 259±47 | 286±36 | H: **0.0022**<br>G: 0.3970<br>H x G: 0.2958 | 0.9739 / 0.3673 |
| Total endpoints<br>(number) | 116±12<br>ns | 119±14<br>ns | 116±12 | 115±13 | H: 0.4002<br>G: 0.8103<br>H x G: 0.1855 | 0.7356 / 0.9624 |
| Mean E Lacunarity | 0.37±0.02<br>*** | 0.36±0.05<br>*** | 0.43±0.04 | 0.42±0.07 | H: **<0.0001**<br>G: 9.604<br>H x G: 0.4832 | 0.8672 / 0.7564 |
| Cortical |  |  |  |  |  |  |
| Junctions density<br>(μm/μm²) | 3E-05±5E-06<br>ns | 3E-05±4E-06<br>ns | 3E-05±6E-06 | 2E-05±3E-06 | H: **0.0137**<br>G: **0.0745**<br>H x G: 0.8251 | 0.1968 / 0.3073 |
| Vessels density<br>(μm/μm²) | 2.30±0.17<br>ns | 2.23±0.17<br>ns | 2.31±0.15 | 2.18±0.18 | H: 0.2147<br>G: 0.5339<br>H x G: 0.8217 | 0.5163 / 0.1486 |
| Total length<br>(μm) | 13339±1026<br>ns | 12696±1403<br>ns | 12969±1003 | 12224±1211 | H: 0.1811<br>G: 0.1125<br>H x G: 0.8663 | 0.3924 / 0.2881 |
| Average vessels length<br>(μm) | 306 (69)<br>* | 312 (95)<br># | 258 (77) | 265 (70) | H: **0.0014**<br>G: 0.9781<br>H x G: 0.6237 | 0.9517 / 0.9664 |
| **Total endpoints**<br>(number) | 126 (27)<br>ns | 124 (12)<br>ns | 139 (20) | 128 (25) | H: 0.0736<br>G: 0.0730<br>H x G: 0.4303 | 0.4717 / 0.0981 |
| **Mean E Lacunarity** | 0.30±0.04<br>ns | 0.31±0.03<br>ns | 0.29±0.03 | 0.32±0.04 | H: 0.6548<br>G: 0.1045<br>H x G: 0.2759 | 0.5239 / 0.0972 |

Data represented as mean±SD or median (IQR) for parametric or non-parametric analyses, respectively. # p<0.10 vs. respective contralateral, \*p<0.05 vs. respective contralateral, \*\*\*p<0.001 vs. respective contralateral, \*\*\*\*p<0.0001 vs. respective contralateral. 2w ANOVA = Two-way ANOVA, REML = Mixed effects model, H = Hemisphere, G = Genotype, H x G = Interaction Hemisphere x Genotype.

## 4. Discussion

Cell-specific IL-1α and IL-1β expression and function remain poorly defined in preclinical stroke models. Previous work by our group demonstrated distinct spatiotemporal expression of IL-1α and IL-1β to suggest non-redundant functions in neuroinflammation post-stroke^5, 6^. Whilst the acute role of IL-1β in promoting neuronal apoptosis and exacerbating ischemic injury has been established^20, 21^, its microglial-specific contribution to both short- and long-term post-stroke cerebrovascular injury and/or repair remains unclear. Here, microglial IL-1β deletion shaped the early inflammatory response to stroke without exerting major long-term effects, pointing to a phase- and niche-specific contribution of IL-1 to cerebrovascular injury and repair.

IL-1 is a pleiotropic cytokine directly participating in the inflammatory response in both infectious and sterile inflammation, activating the innate immune response inducing the production of chemokines, cytokines such as TNFα, IL-6 and IL-1 itself, promoting neutrophilia, but also activating and amplifying the adaptive immune response^22^, and participating in other non-immune biological processes^23, 24^. We recently showed that IL-1α expression peaks acutely and is exclusively expressed by microglia whereas delayed IL-1β expression was observed in a small subset of microglia (11%) and infiltrating neutrophils, in a model of permanent thrombotic ischemic stroke^6^. In this study, we observed that less than 5% of microglia were IL-1β+ at 24 h post-MCAO. To our surprise, a partial deletion without altering IL-1β production by other myeloid populations nor microglial IL-1α levels was sufficient to improve early neurological outcome and modestly limit infarct size and BBB leakage, dampen systemic cytokines and reduce brain neutrophil recruitment, while leaving early immune-cell composition and microglial activation largely unchanged. These findings are consistent with previous studies showing that IL-1 drives ischemic brain injury partly by orchestrating a multicellular cascade that promotes neutrophil recruitment, with γδ T cell-derived IL-17A and astrocyte-derived CXCL1 as key IL-1-responsive intermediates^25^, and brain endothelial IL-1R1 activation promoting BBB disruption and adhesion molecule upregulation^17^. Our findings further indicate that a very small IL-1β⁺ microglial subset can act as an upstream amplifier of the IL-1-driven inflammatory cascade.

At later time points, microglial IL-1β deletion did not alter overall long-term neurological recovery, peri-infarct angiogenesis, pericyte coverage or astrogliosis, in contrast to our previous work showing that microglial IL-1α does not affect acute outcomes nor lesion progression but is required for optimal neurorepair and functional recovery^6^. Despite the contribution of neutrophil extracellular traps (NETs) to reduced post-stroke neovascularization^26^ and considering the acute effect of microglial IL-1β deletion on neutrophil recruitment, we did not observe changes in vessel density, vessel network morphology nor pericyte density and coverage in the peri-infarct regions at 14 days. In line with this, previous studies have shown that IL-1α, but not IL-1β, selectively triggers angiogenesis *in vitro*^27^. However, our results further revealed a striking regional dissociation in neurogenic responses. Deletion of microglial IL-1β enhanced the SVZ neurogenic pool and ectopic SVZ-derived neuroblast migration into the peri-lesional striatum, which is in line with reports that IL-1Ra treatment after stroke promotes long-term neurogenesis and functional recovery^28^. In contrast, microglial IL-1β deletion tended to blunt hippocampal SGZ neurogenesis, without altering neuronal incorporation in the dentate gyrus at 14 days post-stroke. These observations may reflect a region-specific coupling between injury severity and neurogenic response. A plausible interpretation is that smaller early lesions and reduced inflammatory injury in mIL-1β-deficient mice lower the injury-driven proliferative drive in the hippocampus, while preserving or even improving SVZ niche integrity and pro-neurogenic cues in peri-infarct territories. However, microglia are recognized as essential regulators of adult hippocampal neurogenesis, providing trophic factors, clearing apoptotic newborn cells, and shaping circuit integration, which has been shown to partly depend on CX3CL-CX3CR1 signaling, with mice lacking CX3CR1 showing reduced SGZ neurogenesis^29^. In this light, the modest suppression of stroke-induced SGZ neurogenesis observed here could reflect the combined effect of a reduced IL-1β-driven proliferative drive and a baseline vulnerability of the dentate gyrus niche due to CX3CR1 haploinsufficiency, rather than IL-1β loss alone. Nonetheless, previous studies have shown that IL-1β and microglia exert niche- and context-dependent effects on neurogenesis and fate specification^3, 8, 30, 31^, leading to a complex picture in which net effects on neurogenesis may reflect integrated changes in trophic support, the inflammatory milieu and vascular status rather than a single direct action on neural stem cells. Importantly, stroke-induced neurogenesis in the hippocampus can be maladaptive and has been linked to aberrant circuit remodeling and memory deficits^32^. Our results therefore emphasize the need to examine chronic cognitive outcomes and to dissect how microglial IL-1β shapes different neurogenic niches and circuits beyond 2 weeks post-stroke.

We also showed that acutely after ischemic stroke, a much larger fraction of microglia are IL-1α⁺ than IL-1β⁺, which is in line with our previous findings, while double-positive cells are rare, strongly suggesting functional specialization. This raises key unresolved questions, including why microglia preferentially adopt an IL-1α⁺, IL-1β⁻ state; whether IL-1β⁺ microglia represent a distinct transcriptional/proteomic subset or a transient state along an activation trajectory; and how these subsets map onto known disease-associated or neurogenic-niche microglial phenotypes described in stroke and other central nervous system (CNS) injuries. To the best of our knowledge, this is the first study assessing IL-1α and IL-1β expression simultaneously at the single-cell level in microglia after stroke. We nonetheless acknowledge that there are important caveats in our approach. Flow cytometry provides snapshots at discrete timepoints but cannot fully resolve rapid switching between IL-1α⁺ and IL-1β⁺ states and is constrained by antibody sensitivity and intracellular staining conditions. Future studies using dual IL-1α/IL-1β reporter lines to perform longitudinal fate-mapping of IL-1-expressing cells, alongside single-cell transcriptomics and spatial profiling are required to define whether distinct IL-1α+-only, IL-1β+-only and IL-1α/β-double-positive subsets exist, how they evolve over time, and how they relate to local microenvironments such as the peri-infarct region and different neurogenic niches.

Notably, the beneficial effects of microglial IL-1β deletion appeared more pronounced in female mice, with greater improvements in acute neurological outcome, reduced infarct size, and attenuated systemic inflammation. While these observations should be interpreted with caution given the limited statistical power of sex-stratified analyses, these findings align with extensive evidence for sex-dependent regulation of immune responses^33^ and emphasize the importance of further investigation into sex-specific IL-1-dependent mechanisms in ischemic stroke.

It is important to highlight not only the power of the Cx3cr1-CreER–based methodology leveraged here, but to acknowledge its limitations. Cre/loxP-based microglial lines remain the main *in vivo* genetic manipulation strategy due to technical challenges around viral and electroporation-based delivery in microglia, and Tet-On/Off systems requiring the use of immunosuppression. Particularly, CreER lines leveraging the *Cx3cr1* promoter remain the most used method to target microglia which, compared with other lines such as *Sall1*^CreER^ or *Cd11b*^CreER^ show less recombination in other brain or immune cell types^34^. The timing of tamoxifen relative to stroke was chosen to exploit the turnover difference between long-lived microglia and short-lived circulating monocytes which also express CX3CR1^10^, and the functional and flow-cytometry data argue for efficient microglial IL-1β deletion without major off-target effects on other IL-1 family members or basal microglial activation. However, while our model largely targets microglia, we cannot rule out the effects of IL-1β recombination in long-lived CNS BAMs, including perivascular, leptomeningeal and choroid plexus macrophages^35^. Furthermore, the mIL-1β KO condition partially ablates endogenous expression of CX3CR1, which has been implicated in microglial chemotaxis, microglia-mediated neurotoxicity, and microglial activation. However, our first experiments exploring the effect of Cx3cr1^CRE-ERT2^:IL-1β^fl/fl^ recombination on acute stroke outcomes included a non-tamoxifen treated Cx3cr1^CRE-ERT2^:IL-1β^fl/fl^ control group showed no effects on microglial IL-1β deletion, IL-β nor IL-1α expression in other cell types, microglial activation nor acute stroke outcome.

Altogether, our results suggest that IL-1β produced by microglia and potentially other long-lived CX3CR1⁺ CNS BAMs is a key amplifier of early post-stroke injury while having more nuanced, region-specific effects on subacute neurogenesis. These findings build on our previously published work, supporting a model in which endogenous microglial IL-1α and IL-1β serve distinct, non-redundant roles after stroke: IL-1β predominantly drives acute inflammation, BBB disruption and neutrophil-mediated injury, whereas IL-1α contributes to subacute neurorepair and vascular remodeling. It is worth noting that cell-type-specific IL-1R1 studies already indicate that discrete IL-1 signaling modules (endothelial, myeloid, neuronal) serve different functions^3^. Our data suggest that a similar “modular” logic may apply upstream at the level of IL-1α/β-producing microglial subsets, which future reporter-based and conditional KO studies should dissect explicitly. From a translational perspective, early phase stroke trials have mainly shown biological (anti-inflammatory) benefit, and systemic IL-1R1 blockade with anakinra appears mechanistically aligned with the present study. High-dose anakinra regimens are safe, can achieve therapeutically relevant CNS exposure through a disrupted BBB, and improve outcome in preclinical ischemia-reperfusion models and selected patient groups^36–38^. Anakinra crosses the BBB more effectively than large IL-1α/β-neutralizing antibodies and therefore remains the most realistic IL-1-targeting candidate for ischemic stroke, particularly in patients with salvageable tissue, although permanent large-vessel occlusion and malignant infarction may be less amenable to this strategy^39, 40^. At the same time, our findings warn that prolonged, non-selective IL-1 blockade could compromise beneficial neurorepair, arguing for time-limited IL-1R1 antagonism focused on the acute phase. Developing better brain-penetrant, IL-1β-focused or cell-targeted approaches, such as microglia-targeted lipid nanoparticles, could be designed to selectively accumulate and release cargo in response to stroke-related cues, thereby delivering strong but specific and short-lived IL-1β inhibition during the acute damaging window while minimizing interference with later IL-1-dependent neurorepair.

## Acknowledgements

The authors would like to thank the Bioimaging and Flow Cytometry Facilities in the FBMH at the University of Manchester for equipment, training and advice, and Dr Andrew Greenhalgh for advice on flow cytometry analyses.

## Data availability

All data are available upon reasonable request to the corresponding author.

## Authors contributions

A.G.: methodology, experiments execution, data analysis and interpretation, figure creation, writing and editing. M.B.: experiments execution, data analysis and interpretation, and editing. K.W.: experiments execution, data interpretation and editing. R.T.: experiments execution, data analysis and editing. G.C.: experiments execution, data analysis and editing. S.J.: experiments execution, data analysis and editing. N.L.: experiments execution and editing. J.R.C.: experiments execution and editing. J.E.K.: project supervision and editing. D.B.: conceptualization, project supervision and editing. S.M.A.: conceptualization, project supervision and editing. E.P.: conceptualization, project supervision and administration, and editing.

## Ethics approval for the use of animals

All animal procedures were carried out in accordance with the Animals (Scientific Procedures) Act (1986), under a Home Office UK project license, approved by the local Animal Welfare Ethical Review Board, and experiments were performed in accordance with ARRIVE (Animal Research: Reporting of In Vivo Experiments) guidelines, with researchers blinded to experimental groups.

