## Supplementary Information for "Specific deletion of interleukin-1 beta in microglia improves acute outcome and modulates neurogenesis after ischemic stroke"

**Short title**: Microglial interleukin-1 β in ischemic stroke

Alba Grayston[1,2], Margarida Baptista[1,2], Kelly Wemyss[3], Ruby Taylor[1,2], Grace Cullen[1,2], Syeda S Jafree[1,2], Nadim Luka[1,2], Joshua R Cox[3], Joanne E Konkel[3], David Brough[1,2], Stuart M Allan[1,2], Emmanuel Pinteaux[1,2]*.

[1]Division of Neuroscience, School of Biological Sciences, Faculty of Biology, Medicine and Health (FBMH), The University of Manchester, Manchester, UK.

[2]Geoffrey Jefferson Brain Research Centre, University of Manchester, Northern Care Alliance NHS Foundation Trust, The Manchester Academic Health Science Centre, Manchester, UK.

[3]Lydia Becker Institute of Immunology and Inflammation, School of Biological Sciences, Faculty of Biology, Medicine and Health, Manchester Academic Health Science Centre, University of Manchester, Manchester, UK.

#### Tables

**Table S1.** **Plasma cytokine levels at 24 h after MCAO**

| Cytokines | WT (pg/mL) | WT-Cx3cr1-Cre (pg/mL) | mIL-1β KO (pg/mL) | ANOVA/K-W (p-value) |
|---|---|---|---|---|
| **IL-23** | 380.7 (0) | 432.7 (93.1) | 531.5 (0) | 0.5000 |
| **IL-1α** | 17.31 (23.1) | 5.54 (3.2) | 27.68 (41.2) | 0.3619 |
| **TNFα** | 103.3 (115.1) | 185.8 (220.3) | 57.4 (42.4) | 0.3619 |
| **IFNγ** | 10.9 (1.2) | 15.7 (4.3) | 22.5 (14.3) | 0.0929 |
| **MCP-1** | 182.9 (0) | 119.1 (178.1) | 63.0 (548.9) | 0.9000 |
| **IL-12p70** | 15.8 (4.9) | 15.5 (1.0) | 14.7 (17.4) | >0.9999 |
| **IL-1β** | 58.8 (52.3) | 48.1 (41.2) | 89.9 (85.5) | 0.0857 |
| **IL-10** | 196.8 (102.1) | 227.8 (549.9) | 119.3 (439) | 0.5016 |
| **IL-6** | 151.1 (127.9) | 54.73 (1.0) | 61.1 (58.6) | 0.7571 |
| **IL-27** | 631.8 (257.7) | 551.8 (724.6) | 492.5 (1859) | 0.8444 |
| **IL-17A** | 6.4 (2.6) | 8.1 (27.9) | 14.9 (19.6) | 0.3762 |
| **IFN-β** | 269.6 (0) | 389.1 (350.6) | 290.9 (729.9) | 0.9571 |
| **GM-CSF** | 27.8 (25.3) | 29.7 (49.3) | 25.5 (19.5) | 0.6730 |

**Table S2.** **Plasma cytokine levels at 14 days after MCAO**

| Cytokines | WT (pg/mL) | mIL-1β KO (pg/mL) | t-test/M-W (p-value) |
|---|---|---|---|
| **IL-23** | 746 (1013) | 860.8 (1834) | 0.7214 |
| **IL-1α** | 7.06 (24.8) | 8.97 (15.5) | 0.5783 |
| **TNFα** | 183.3 (470.8) | 168.6 (268.2) | 0.9803 |
| **IFNγ** | 41.8 (120.1) | 46.6 (111.0) | 0.8741 |
| **MCP-1** | 35.4 (42.1) | 48.3 (72.5) | 0.1859 |
| **IL-12p70** | 58.6 (148.6) | 50.0 (77.4) | 0.2872 |
| **IL-1β** | 59.3 (162.6) | 59.6 (78.1) | 0.7863 |
| **IL-10** | 345.8 (692.7) | 240.2 (469.2) | 0.1912 |
| **IL-6** | 40.4 (90.6) | 28.3 (68.4) | 0.4063 |
| **IL-27** | 1487 (3930) | 1363 (2519) | 0.8172 |
| **IL-17A** | 41.0 (84.4) | 26.6 (54.1) | 0.2409 |
| **IFN-β** | 731.7 (1755) | 536.2 (1713) | 0.8042 |
| **GM-CSF** | 64.1 (113) | 66.9 (73.3) | 0.6340 |

### Figures

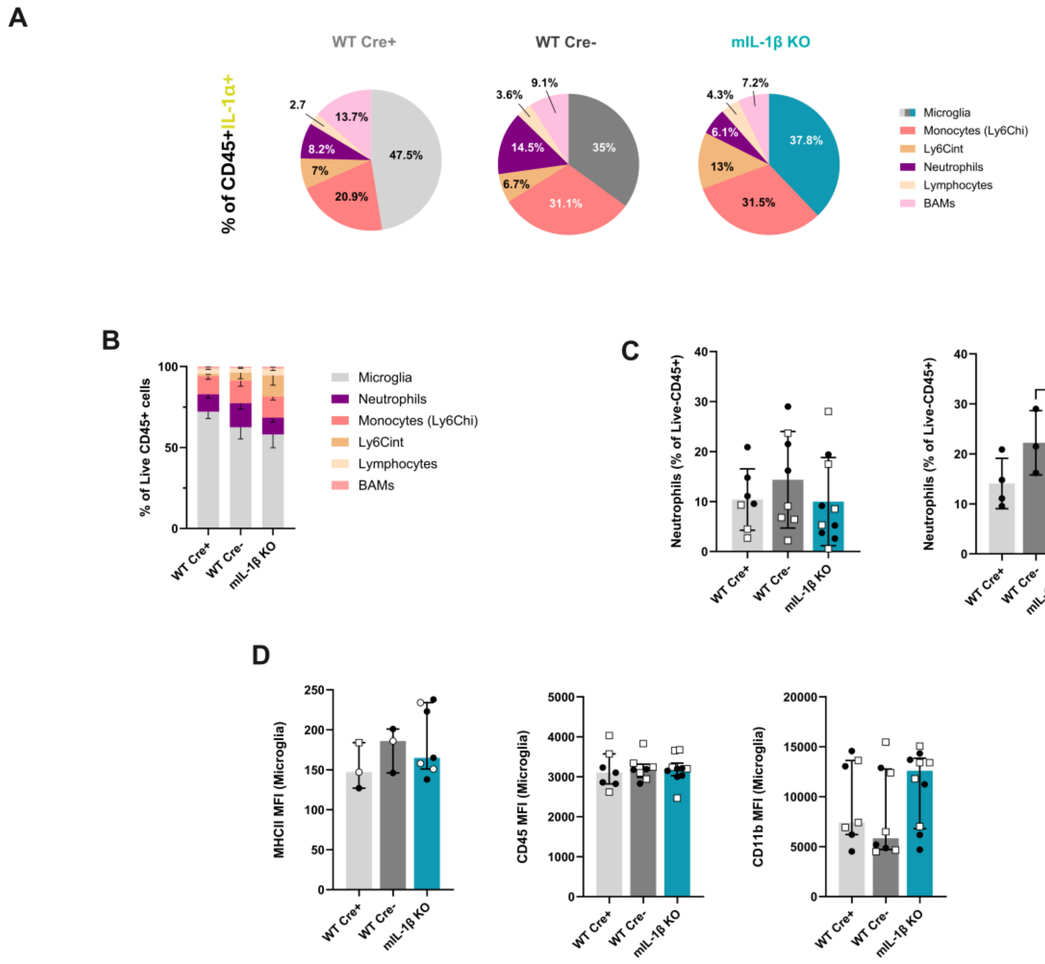


**Figure S1. Microglial IL-1β deletion does not affect the immune cell landscape in the brain at 24 h post-MCAO. A)** Representative gate of IL-1α+ cells within the Live CD45+ cell population (left) and pie charts (left) showing the distribution of IL-1α+ immune cells, shown as the mean percentage of cell populations in ipsilateral brain samples across genotypes. **B)** Relative distribution of immune cell types within the Live CD45+ cell population in the ipsilateral brain across genotypes. **C)** Proportion of neutrophils, expressed as the percentage of Live CD45+ cells, in both males and females (left) and in only females (right). **D)** Graphs showing the mean fluorescence intensity (MFI) of microglial MHC II, CD45 and CD11b. *p<0.05

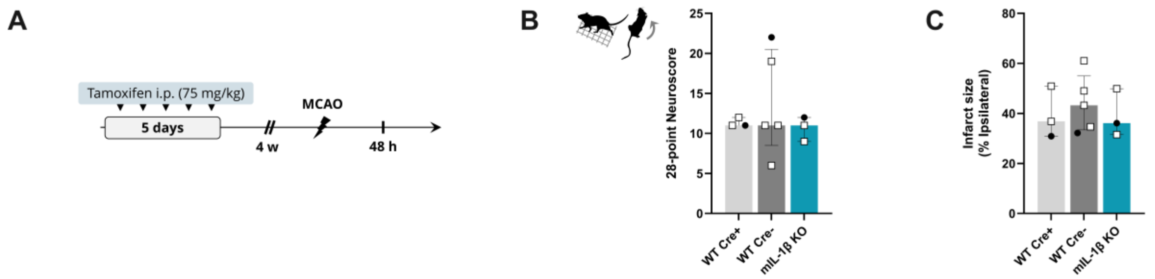


**Figure S2. CX3CR1 haploinsufficiency does not modify acute stroke outcome. A)** Schematic representation of the experimental design. **B)** Neurological outcome at 48 h post-MCAO assessed by a 28-point based neurological score (n=3-5, Kruskal-Wallis test). **C)** Infarct size quantification expressed as the percentage of total ipsilateral hemisphere volume (n=3-4, Kruskal-Wallis test).

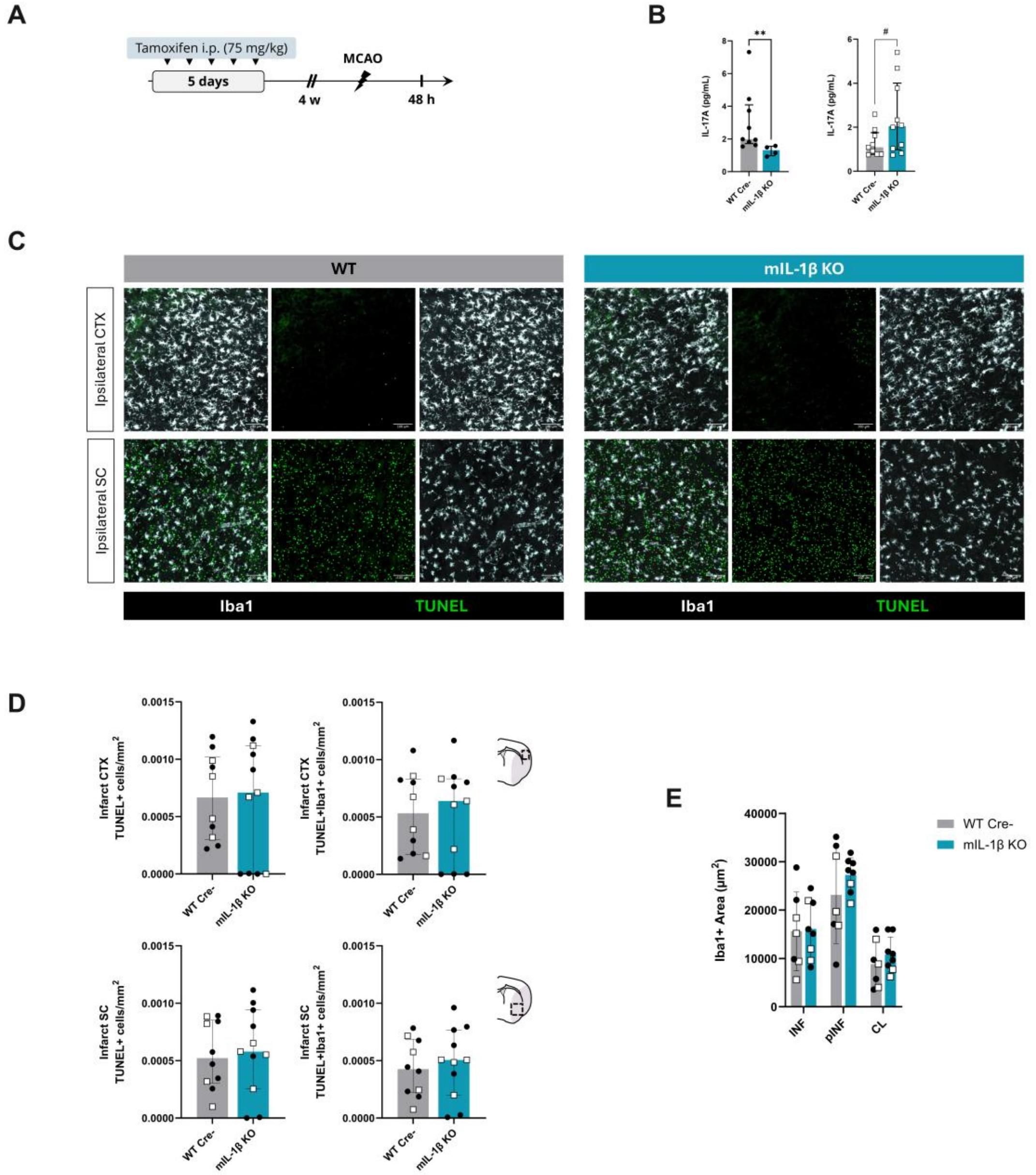

**Figure S3. Microglial IL-1β deletion shows sex differences in IL-17A expression and does not affect cell death nor microglial activation at 2 days post-MCAO. A)** Schematic representation of the experimental design. **B)** Plasma IL-17A levels in females (left) and in males (right). **C)** Representative TUNEL staining images in the cortical (top) and subcortical infarct regions (bottom). **D)** TUNEL+ cells density in the cortical (top) and subcortical infarct regions (bottom). **E)** Microglial (Iba1+) area in the infarct, peri-infarct and healthy contralateral brain regions. INF: infarct, pINF: peri-infarct, CL: contralateral.

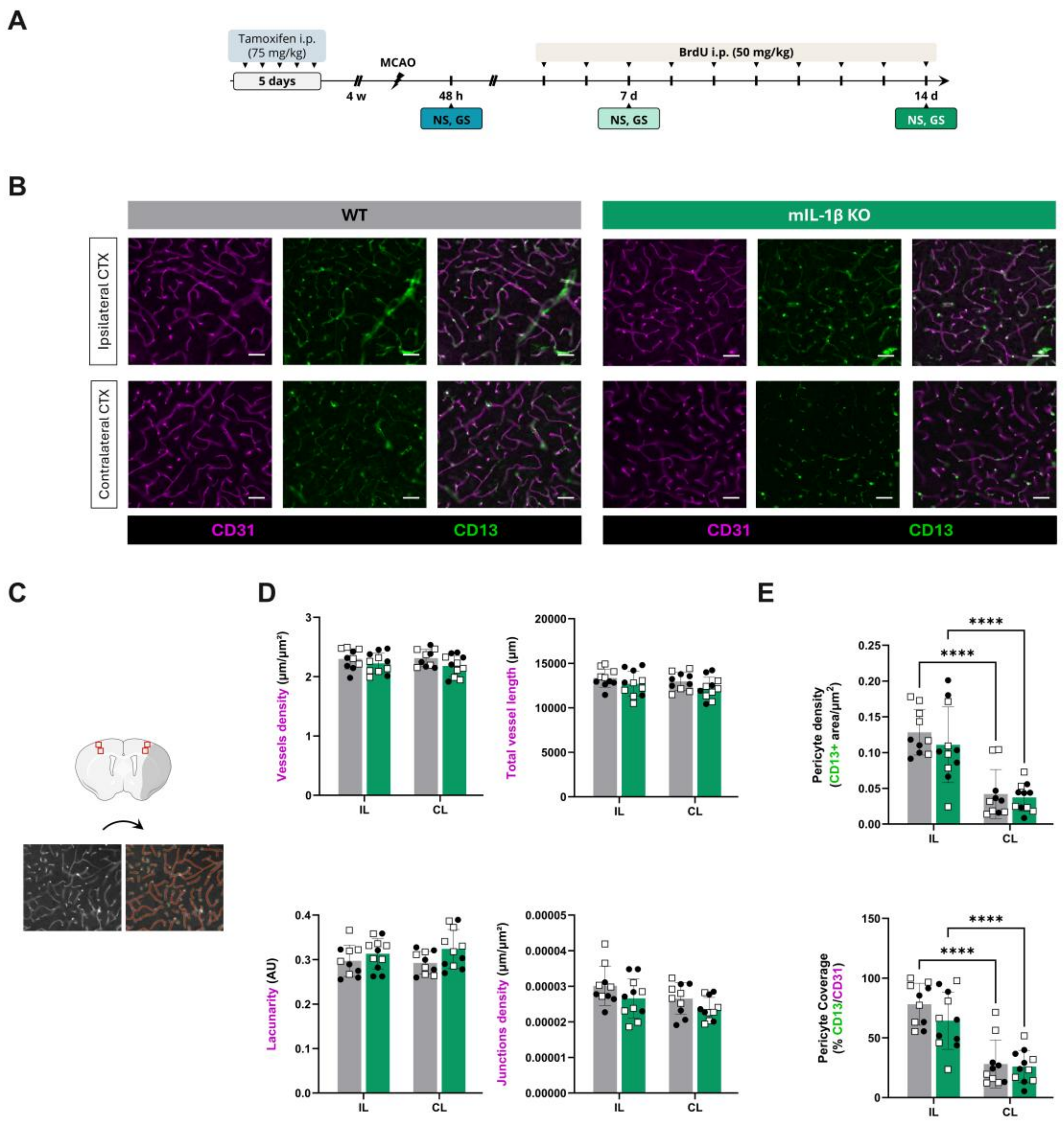


**Figure S4. Microglial IL-1β deletion does not affect post-stroke angiogenesis. A)** Schematic representation of the experimental design. **B)** Representative immunostaining of the vasculature, stained for CD31 (endothelial cells) and CD13 (pericytes) in the peri-infarct subcortical region (SC) in the ipsilateral (IL) hemisphere and reflective contralateral (CL) cortical region of WT and mIL-1β KO mice (scale bar: 50 μm). **C)** Schematic of a coronal brain section at bregma +0.3 showing ROIs selected for analyses of the cortical areas, depicting the infarct region in grey (top) and representative CD31-stain image for vessel network analysis with AngioTool (left) and output image (right) showing the outline of the vasculature (yellow), skeleton representation of the vasculature (red) and branching points (blue). **D)** Graphs presenting selected vessel network analysis outputs, including vessels density, total vessel length, lacunarity and junctions density (n=10-11/group, 2-way ANOVA, and REML for junctions density analysis, followed by Šídák's multiple comparisons test). **E)** Quantification of pericyte (CD13+) cell density (top) and pericyte coverage, expressed as the percentage of CD13+ area per CD31+ vessels area, in the IL and CL peri-infarct cortical areas (n=8-12/group, 2-way ANOVA followed by Šídák's multiple comparisons test, ****p<0.0001). CTX: cortex.
